# Linkage Disequilibrium Estimation in Low Coverage High-Throughput Sequencing Data

**DOI:** 10.1101/235937

**Authors:** Timothy P. Bilton, John C. McEwan, Shannon M. Clarke, Rudiger Brauning, Tracey C. van Stijn, Suzanne J. Rowe, Ken G. Dodds

## Abstract

High-throughput sequencing methods that multiplex a large number of individuals have provided a cost-effective approach for discovering genome-wide genetic variation in large populations. These sequencing methods are increasingly being utilized in population genetic studies across a diverse range of species. One side-effect of these methods, however, is that one or more alleles at a particular locus may not be sequenced, particularly when the sequencing depth is low, resulting in some heterozygous genotypes being called as homozygous. Under-called heterozygous genotypes have a profound effect on the estimation of linkage disequilibrium and, if not taken into account, leads to inaccurate estimates. We developed a new likelihood method, GUS-LD, to estimate pairwise linkage disequilibrium using low coverage sequencing data that accounts for under-called heterozygous genotypes. Our findings show that accurate estimates were obtained using GUS-LD on low coverage sequencing data, whereas underestimation of linkage disequilibrium results if no adjustment is made for under-called heterozygotes.

---

LINKAGE disequilibrium (LD) is the term given to the non-random association of alleles located at different loci in a population. Quantifying the level of LD or estimating the pairwise LD between all loci in a population is of interest to many researchers as it has many important applications. For example, in association mapping studies, LD is used to identify candidate regions of the genome associated with a particular trait or disease and can provide finer resolution in mapping compared to linkage based studies (Devlin and Risch 1995; Jorde 1995; Xiong and Guo 1997; Mackay and Powell 2007). LD is affected by many genetic and evolutionary forces such as recombination, admixture, migration, selection and gene flow among others (Terwilliger *et al.* 1998; Ardlie *et al.* 2002; Gaut and Long 2003; Slatkin 2008). Consequently, LD patterns can be used to quantify genetic diversity and make inferences about the evolutionary history of natural populations (Nordborg and Tavaré 2002; Slatkin 2008; Zhu *et al.* 2015). In addition, the relationship between map distance and the level of LD can be used to estimate the effective population size (Hill 1981; Hayes *et al.* 2003; Waples 2006; Sved *et al.* 2013).

Today, many species are being sequenced using high-throughput sequencing methods that multiplex a large number of individuals. Some of the most popular sequencing methods are whole genome sequencing and the reduced representation approaches such as genotyping-by-sequencing (Elshire *et al.* 2011), whole-exome sequencing (Hodges *et al.* 2007), and restriction-site associated DNA (Baird *et al.* 2008). These sequencing methods provide a low cost approach for performing genome-wide genotyping and discovery of single nucleotide polymorphisms (SNPs). Genetic data generated using these sequencing methods are increasingly being used to compute pairwise linkage disequilibrium estimates (e.g., Hohenlohe *et al.* (2012); Wang *et al.* (2013); Nimmakayala *et al.* (2014); Huang *et al.* (2014); Xu *et al.* (2014); Fè *et al.* (2015); Zhang *et al.* (2015); Covarrubias-Pazaran *et al.* (2016); Van Wyngaarden *et al.* (2016)). A major disadvantage with these methods, however, is that one or both of the alleles at a particular locus may be missed for a given individual if the sequencing depth is low. If neither allele is seen, a missing genotype results while if only one of the two parental alleles is seen (possibly multiple times), a heterozygous genotype may be called as homozygous (Dodds *et al.* 2015; Fragoso *et al.* 2016). The latter case is also known as allelic dropout and is particularly problematic as genotype calls with this type of missingness behave like genotyping errors, which have a profound impact on the estimation of LD even when the error rates are low (Akey *et al.* 2001).

One way of removing genotyping errors resulting from low sequencing depth is to set genotype calls with an associated read depth below some threshold value to missing. However, such filtering results in fewer individuals and SNPs for a given sequencing cost (Dodds *et al.* 2015), and for low coverage data may result in insufficient data to undertake the analysis. LD is often estimated using haplotypes phased from genotype data via various software packages and algorithms such as BEAGLE (Browning and Browning 2007), fastPHASE (Scheet and Stephens 2006), MaCH (Li *et al.* 2010), and FILLIN (Swarts *et al.* 2014). However, all of these approaches require that the chromosomal order of the loci is known in order to infer the haplotypes, which is not necessarily the case for reduced representation sequencing data, particularly if the SNPs are called *de novo*. A few alternative approaches for estimating LD from high-throughput sequencing data have been presented in the literature. Feder *et al.* (2012) proposed estimating pairwise LD using reads that cover both loci while estimating the allele frequencies using all the reads. This approach, however, is not applicable to short read sequencing data (e.g., genotyping-by-sequencing) where most of the reads do not cover both sites. Alternatively, it restricts the analysis to loci that are very close, which may not be that useful. Maruki and Lynch (2014) presented a likelihood method for estimating the disequilibrium coefficient in situations where there is a combination of reads that intersect both loci or only one of the two loci. Their method also accounts for sequencing errors, that is errors caused by a base being miscalled, but does not account for genotyping errors resulting from under-called heterozygotes.

We present a new method for estimating pairwise LD using low coverage sequencing data, without requiring haplotype phasing, a known chromosomal order or filtering with regard to read depth. In essence, our method is based on the likelihood method by Hill (1974), which estimates LD using genotypic data in random mating populations, but is extended to account for under-called heterozygous genotypes. We also examine the effect genotyping errors from low read depths have on the estimation of LD.

## Materials and Methods

### Estimation of pairwise LD

Let *A_j_* and *B_j_* denote the major and minor allele at locus *j* respectively and let *p_A_j__* and *p_B_j__* denote the major and minor allele frequencies for locus *j* respectively. The linkage disequilibrium coefficient is defined as (Lewontin and Kojima 1960)

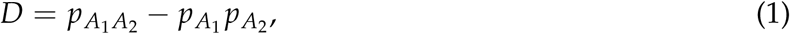

where *p*_*A*_1_*A*_2__ is the probability of observing a haplotype containing the major allele at both loci. Since probabilities are required to be non-negative, *D* must satisfy the constraints (Lewontin 1964)

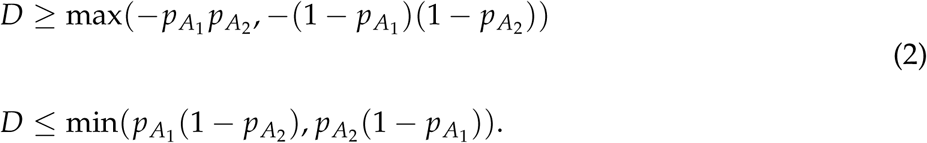

We let *G_ij_* denote the true genotype and 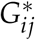 denote the genotype observed in the sequencing data for individual *i* at locus *j* for *i* = 1,…, *n*. The observed genotypes, 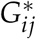, are determined based on the number of unique alleles observed (e.g., 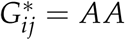 if only the allele *A* is observed regardless of the number of reads sequenced). We let *G_i_* = (*G_ij_, G_ik_*) denote the true joint genotype and 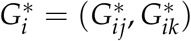 the joint genotype observed in the sequencing data for individual *i* between locus *j* and *k*, where *j* ≠ *k* and *i* = 1,…, *n*. Furthermore, we let *AA_j_, AB_j_,* and *BB_j_* denote the major homozygous genotype, heterozygous genotype, and minor homozygous genotype at locus *j* respectively. For two biallelic loci, the nine joint genotypes are (*AA*_1_, *AA*_2_), (*AA*_1_, *AB*_2_), (*AA*_1_, *BB*_2_), (*AB*_1_, *AA*_2_), (*AB*_1_, *AB*_2_), (*AB*_1_, *BB*_2_), (*BB*1, *AA*_2_), (*BB*_1_, *AB*_2_), and (*BB*_1_, *BB*_2_), which we denote by 1 to 9 respectively. By the law of total probability,

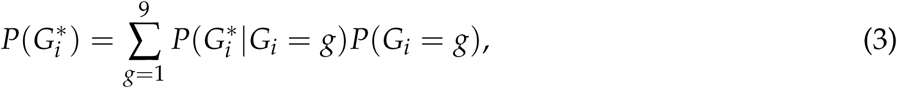

where *g* = (*g_1_,g_2_*). Under the assumption that the probabilities of observing a particular genotype in the sequencing data given the true genotype are independent between loci, expression (3) simplifies to,

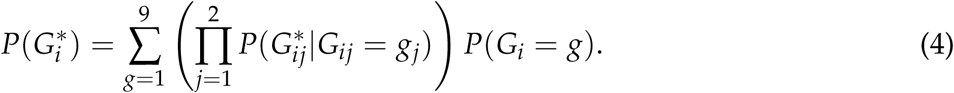

The expected true joint genotype probabilities, *p_ig_* = *P*(*G_i_* = *g*), correspond to those given in Table 1 when the population is in Hardy-Weinberg equilibrium (Hill 1974).

**Table 1.** Joint genotype probabilities for two biallelic loci under the assumption of Hardy-Weinberg equilibrium

| $g$ | Locus 1 | Locus 2 | $p_{ig}$ |
| --- | --- | --- | --- |
| 1 | $AA_1$ | $AA_2$ | $(p_{A_1}p_{A_2} + D)^2$ |
| 2 | | $AB_2$ | $2(p_{A_1}p_{A_2} + D)(p_{A_1}p_{B_2} - D)$ |
| 3 | | $BB_2$ | $(p_{A_1}p_{B_2} - D)^2$ |
| 4 | $AB_1$ | $AA_2$ | $2(p_{A_1}p_{B_2} + D)(p_{B_1}p_{A_2} - D)$ |
| 5 | | $AB_2$ | $2(p_{A_1}p_{A_2} + D)(p_{B_1}p_{B_2} + D) + 2(p_{A_1}p_{B_2} + D)(p_{B_1}p_{A_2} - D)$ |
| 6 | | $BB_2$ | $2(p_{A_1}p_{B_2} + D)(p_{B_1}p_{B_2} + D)$ |
| 7 | $BB_1$ | $AA_2$ | $(p_{B_1}p_{A_2} + D)^2$ |
| 8 | | $AB_2$ | $2(p_{B_1}p_{A_2} + D)(p_{B_1}p_{B_2} - D)$ |
| 9 | | $BB_2$ | $(p_{B_1}p_{B_2} - D)^2$ |

The genotypes observed in the sequencing data, 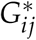, can be considered as arising from a sample of the two alleles found in the true genotype *G_ij_*. If the alleles are read at random and there is no sequencing error present, the conditional probabilities of observing a joint genotype in the sequencing data given the true joint genotype are,

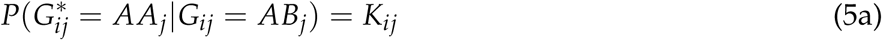

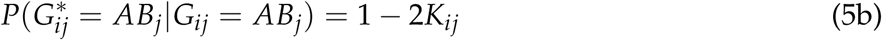

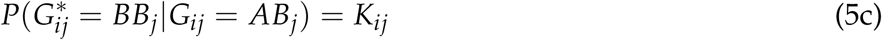

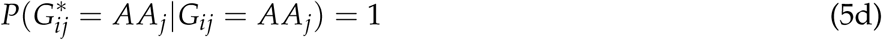

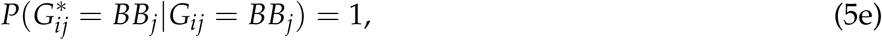

where *K_ij_* = 1/2 *^k_ij_^* and *k_ij_* is the sequencing depth in individual *i* at locus *j* (Dodds *et al.* 2015). The sum of Equations(5a) and (5c) gives the probability of a genotyping error due to low sequencing depth (e.g., a heterozygote being miscalled as homozygous). From Equations(4)-Equations(5e), the joint genotype probabilities for sequencing data, 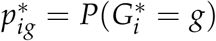, are derived as

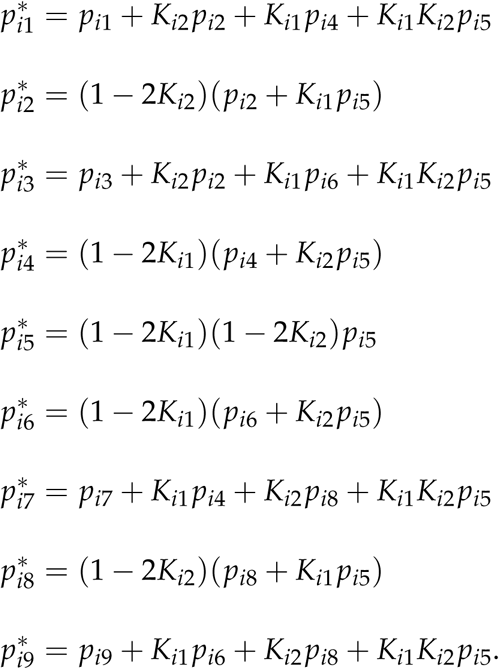

We let 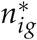 denote the indicator variable of whether the joint genotype, 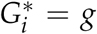, is observed in the sequencing data for individual *i*, where *g* = 1,…, 9 and *i* = 1,…, *n*. Assuming that the distribution of the joint genotype counts for individual *i* follows a multinomial distribution with one trial and probabilities 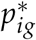, and assuming that the individuals form an independent sample, the log-likelihood for the joint genotype counts observed in the sequencing data is

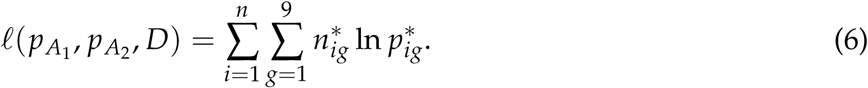

The maximum likelihood estimate of the disequilibrium coefficient, 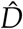, using sequencing data is obtained by maximizing likelihood (6) subject to constraint (2). As no analytical solution exists, maximization of likelihood (6) is performed using numerical methods. The expectation of the maximum likelihood estimate is (Weir 1996),

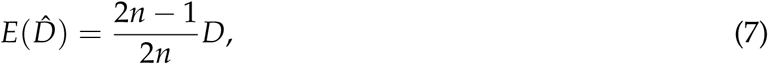

resulting in a small bias which is removed by multiplying 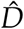 by 2*n*/(2*n* – 1) subject to constraint (2). If there are missing genotypes, *n* is taken as the number of individuals with non-missing genotypes at both loci.

Since the range of *D* depends on the allele frequencies, comparing levels of LD between markers can be difficult using the disequilibrium coefficient. Consequently, many alternative measures of LD have been proposed in the literature; see Hedrick (1987) and Devlin and Risch (1995) for a summary and comparison of these measures. In this article, we shall only consider two commonly used measures, *D*^′^ (Lewontin 1964; Hedrick 1987) and *r*^2^ (Hill and Robertson 1968). Although both *D*^′^ and *r*^2^ are measures of LD, they have different properties and are useful for different applications (see Mueller (2004)). The maximum likelihood estimates for both of these measures are computed using the functions 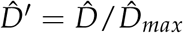 and 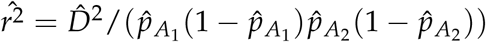, where

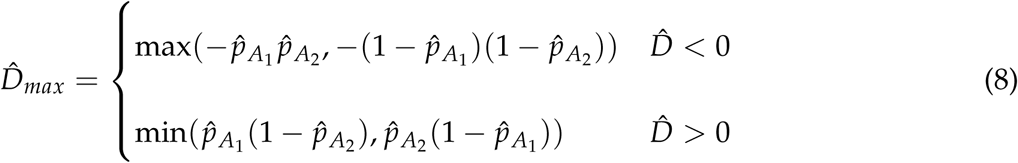

and 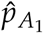 and 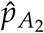 are the maximum likelihood estimates of the major allele frequencies at locus 1 and 2 respectively. We refer to the proposed methodology, which estimates LD adjusting for genotyping errors associated with low sequencing depth, as <u>g</u>enotyping <u>u</u>ncertainty with <u>s</u>equencing data - <u>l</u>inkage <u>d</u>isequilibrium (GUS-LD, pronounced *guzzled*).

### Simulation

To examine the performance of GUS-LD, a simulation study was undertaken. Generation of simulated sequencing data proceeded as follows. For each individual, two haplotypes were sampled from the four possible haplotypes for preset values of *p*_*A*_1__, *p*_*A*_2__ and *D*, and were then converted to genotype calls. Simulation of sequencing data proceeded by firstly generating a read depth for each individual at each locus by simulating realizations from a Poisson distribution with mean *μ_k_j__*, where a range of read depths were used (*μ_k_j__* = 1,2,3,4,5,7.5,10,15). At each locus within each individual, alleles were sampled from the genotype call with equal probability and replacement until a sample size corresponding to the read depth was obtained. In some cases, the simulated read depth was zero resulting in missing genotypes for the simulated sequencing data. The simulations were performed under various combinations of *p*_*A*_1__, *p*_*A*_2__ and *D* (see Table 2 for a list of combinations used).

**Table 2.** Combinations of parameters used in the simulations

| Simulation | $p_{A_1}$ | $p_{A_2}$ | $D$ |
| --- | --- | --- | --- |
| 1 | 0.5 | 0.5 | $-0.15, 0, 0.05, 0.15, 0.25$ |
| 2 | 0.5 | 0.75 | $-0.01, 0, 0.05, 0.1, 0.125$ |
| 3 | 0.9 | 0.9 | $-0.01, 0.03, 0.06, 0.09$ |

Two sets of simulations were performed. The first compares estimation of LD using simulated sequencing data between GUS-LD and the standard likelihood procedure of Hill (1974) that assumes accurate genotype calls. For each combination of parameters, 10,000 simulated datasets of 100 individuals were generated, where estimates of the bias and standard errors of 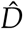, 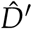 and 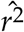 were computed for both methods. In the second set, the optimal sequencing depth for a given sequencing effort, defined as the number of reads which is the product of the number of individuals, the number of loci and the mean read depth, is examined. For each combination of parameters, 10,000 datasets were simulated, where the number of individuals in the datasets were set such that an average sequencing effort of 600 reads was maintained. Estimates of the LD measures were obtained using GUS-LD and the mean square errors of 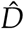, 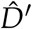 and 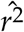 were computed.

### Deer dataset

GUS-LD was also compared to the standard likelihood approach using a dataset consisting of 666 farmed deer and 38 of their sires. The dams were unrecorded Red deer (*Cervus elaphus*) while the sires were predominantly Wapiti (also known as Elk; *Cervus canadensis*), but included some Red deer. The animals were managed in accordance with the provisions of the New Zealand Animal Welfare Act 1999, and the Codes of Welfare developed under sections 68-79 of the Act. Tissue samples were collected in the form of ear tissue punches and DNA extracted according to Clarke *et al.* (2014). Genotyping was performed using the genotyping-by-sequencing method (Elshire *et al.* 2011) using the restriction enzyme *Pst*I and variations of the standard lab methodology as outlined in Dodds *et al.* (2015). The individuals were sequenced across eight lanes at AgResearch, Invermay, Animal Genomics laboratory on an Illimina HiSeq 2500 v4 chemistry yielding approximately 1.34B reads (read length of 1× 100bp) in total. SNP variants were called using UNEAK (Lu *et al.* 2013) as outlined in Dodds *et al.* (2015). For the LD analysis, a set of 38 SNPs that were determined to be close to the microsatellite TGLA94 (Marshall *et al.* 1998), had a minor allele frequency greater than 0.05 and had less than 25% missing genotype calls were retained for analysis.

### Data availability

Scripts for generating the simulated sequencing data are provided in File S1. The deer dataset and an implementation of GUS-LD can be found at https://github.com/AgResearch/GUS-LD.

## Results

### Simulation

For the first set of simulations, the bias for the estimates of the various pairwise LD measures are given in Figure 1, for *p*_*A*1_, = 0.5, *p*_*A*2_ = 0.5 and for a range of values of *D*. When the average read depth was low, the estimates of *D* obtained using the standard likelihood procedure were biased towards zero, where the level of bias increased as the strength of LD increased. In contrast, the estimates computed using GUS-LD were relatively unbiased across the various read depths. Nevertheless, for the cases when *D* was close to or on its upper or lower bound (Equations(2)), 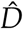 was biased, although the level of bias was much less for GUS-LD than for the standard likelihood procedure. These conclusions, in general, also applied to estimation of *D*^′^ and *r*^2^, although there was some bias in the estimates of *D*^′^ even when the read depth was large and the true value of D was not near the upper or lower bound of its parameter space. This bias is due to poor sampling properties of *D*^′^ and has been observed to occur in simulation studies for small sample sizes (Teare et al. 2002; Terwilliger et al. 2002). As the average read depth increased, the number of under-called heterozygous genotypes in the datasets decreased, which resulted in less bias for the LD estimates obtained from the standard likelihood method. At mean read depths of 10 or more, there were very few under-called heterozygous genotypes present under the simulation model, so that the estimates from the two approaches coincided.

**Figure 1.**
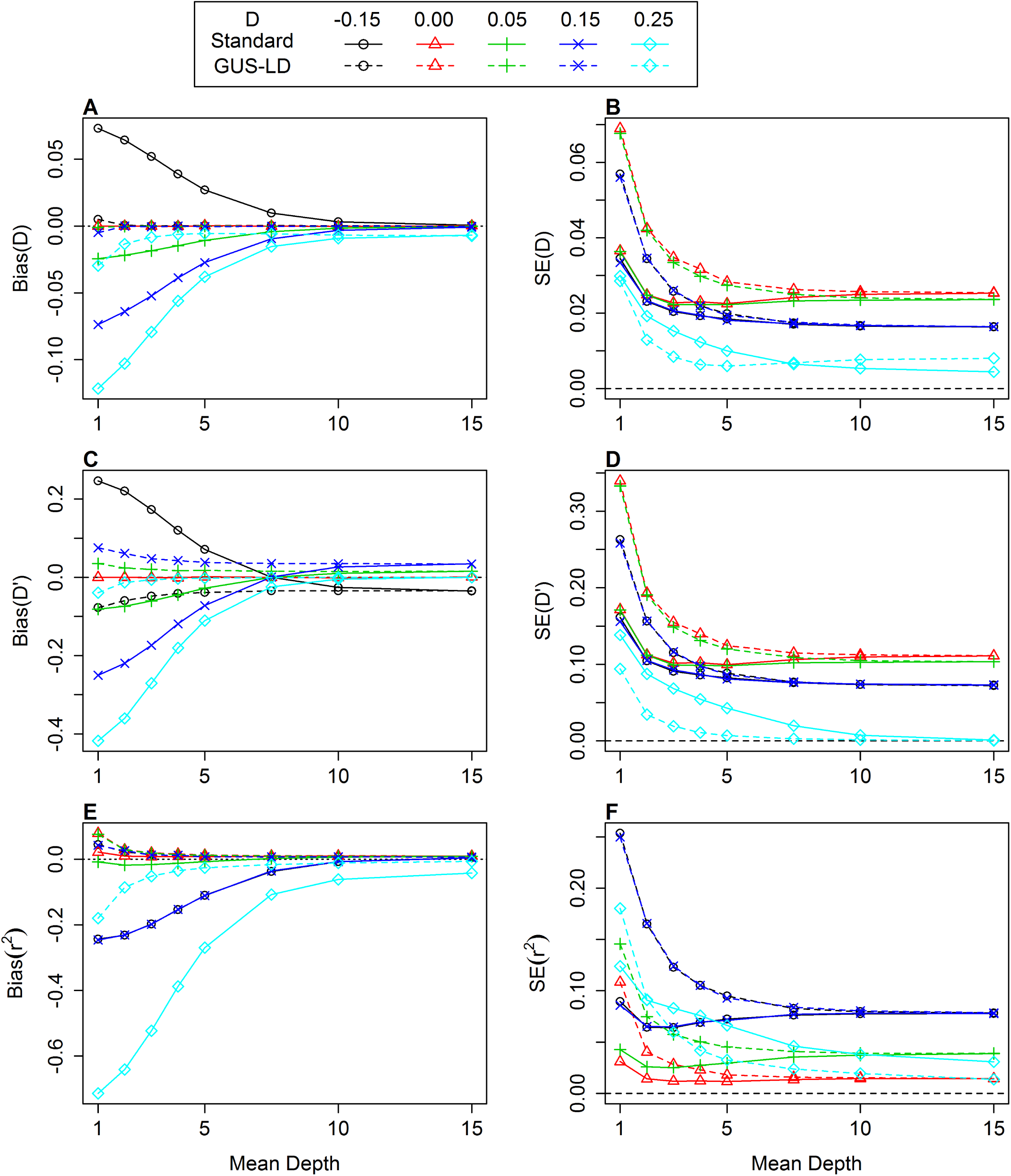
Bias of the estimates for LD measures *D*(A), *D*^′^ (C) and *r*^2^ (E) and standard error (SE) of the estimates for LD measures *D* (B), *D*^′^ (D) and *r*^2^ (F) when *p*_*A*1_ = 0.5 and *p*_*A*2_ = 0.5 and the true values of *D* were –0.05, 0, 0.05, 0.15, and 0.25. The dashed lines represents the estimates obtained using GUS-LD whereas the solid lines represents the estimates obtained using the standard likelihood approach. The upper and lower bound for *D* are –0.25 and 0.25 respectively.

Figure 1 also shows the standard errors of the estimates for the three LD measures computed using the two approaches. In general, the standard errors of the LD estimates computed under GUS-LD were larger compared with those obtained under the standard likelihood approach, with the difference decreasing as the average read depth increased. This increase in the standard errors for GUS-LD was expected as there is extra sampling variation introduced into the sequencing data, caused by not all alleles being observed. On the other hand, when the true value of *D* was close to or on the lower or upper bound of its parameter space (Equations(2)), GUS-LD yielded smaller standard errors than the standard approach for low read depths.

The bias and standard errors of the LD estimates for alternative combinations of allele frequencies are given in Figure S1 *p*_*A*_1__ = 0.5, *p*_*A*_2__ = 0.75) and in Figure S2 *p*_*A*_1__ = *p*_*A*_2__ = 0.9). The results from these simulations were mostly in agreement with those when *p*_*A*_1__ = 0.5 and *p*_*A*_2__ = 0.5.

For the second set of simulations, the mean square error for the estimates of the various pairwise LD measures are given in Figure 2 for a fixed average sequencing effort of 600 reads, *p*_*A*_1__ = *p*_*A*_2__ = 0.5, and for a range of values of *D*. Overall, the mean read depth which gave the lowest mean square error was between 2 and 5, where the actual depth at which the minimum occurs depended on the true value of *D* and the LD measure. There was one exception to this trend that occurred when the true value of D was equal to its upper bound (*D* = 0.25) for the LD measures *D*^′^ and *r*^2^. In this case, the mean square error was largest at smaller mean read depths and decreased as the mean read depth increased. This is due to the fact that there is no variation or bias when the genotypes are accurate for values of *D* that are on its upper or lower bound, but there is variation when there is uncertainty in the genotype calls associated with low read depths.

**Figure 2.**
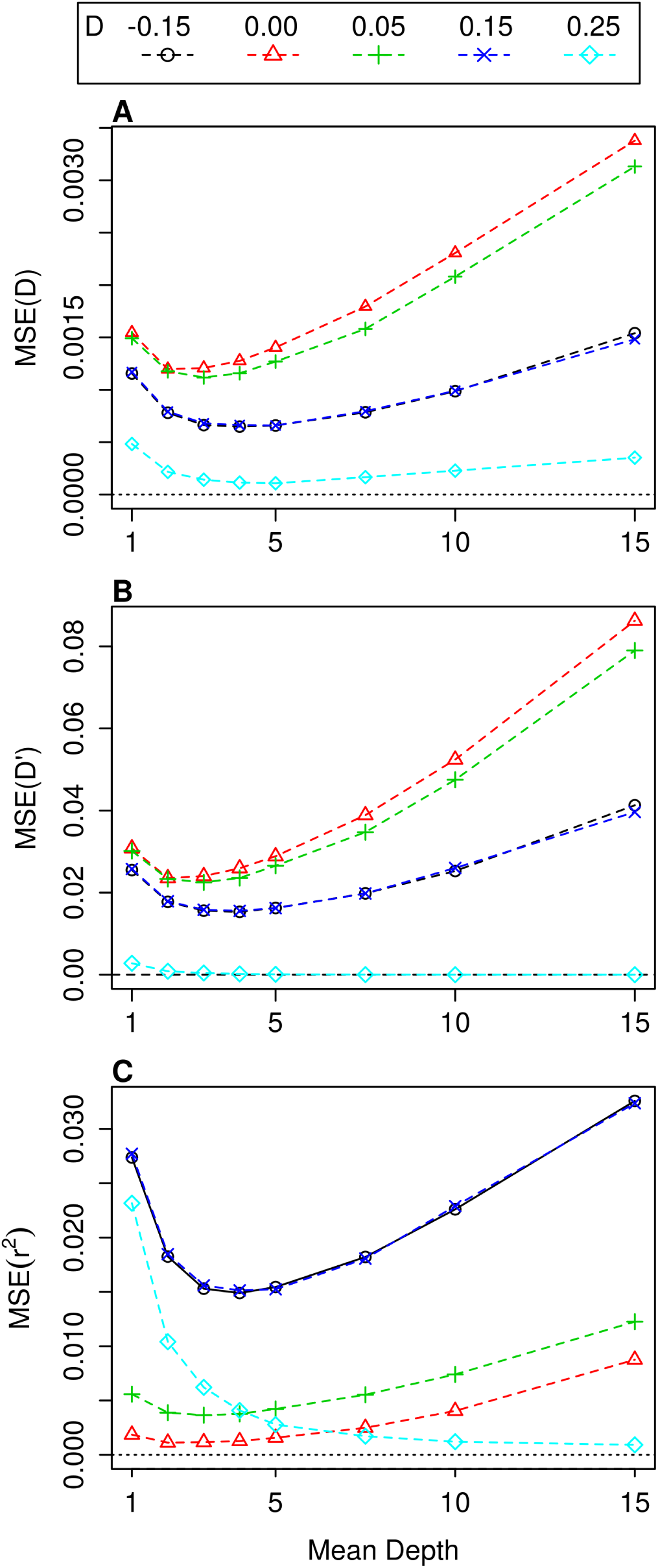
Mean square error (MSE) of the LD measures *D* (A), *D* ^′^ (B) and *r*^2^ (C) for a fixed average sequencing effort of 600 reads when *p*_*A*_1__ = 0.5 and *p*_*A*_2__= 0.5 and the true values of *D* were –0.05, 0, 0.05, 0.15, and 0.25. The LD estimates were obtained using GUS-LD and the upper and lower bound for *D* are –0.25 and 0.25 respectively.

The mean square errors of the LD estimates for alternative combinations of allele frequencies when the sequencing effort was fixed are given in Figure S3 *p*_*A*_1__ = 0.5, *p*_*A*_2__ = 0.75) and in Figure S4 *p*_*A*_1__ = *p*_*A*_2__ = 0.9). The results from these simulations were very similar to the case when *p*_*A*_1__ = *p*_*A*_2__ = 0.5, although there were some differences. For example, the mean square error across all the mean depths for *D* was larger as the true value of *D* increased when *p*_*A*_1__ = *p*_*A*_2__ = 0.9, whereas the reverse was true when *p*_*A*_1__ = *p*_*A*_2__ = 0.5 and when *p*_*A*_1__ = 0.5 and *p*_*A*_2__ = 0.75 Also, for *p*_*A*_1__ = 0.5 and *p*_*A*_2__ = 0.75, the mean square error for the LD measure *r*^2^ was not decreasing as the read depth increased when the true value of *D* was on its upper boundary (*D* = 0.125), as for the other parameter combinations.

This was due to unequal allele frequencies meaning that the estimates of *r*^2^ were not near its upper bound of 1. These differences are due to the complex sampling properties of the various LD measures. Nevertheless, the optimal sequencing depth was mostly between 2 and 5 across all three parameter combinations and LD measures.

### Deer dataset

The LD estimates between all pairs amongst a set of 38 SNPs are given in Figure 3 for the absolute value of *D*^′^ and Figure 4 for *r*^2^. For the former LD measure, a number of pairwise estimates computed using GUS-LD were larger compared to the estimates obtained from the standard likelihood approach, which is seen by the greater intensity of red across the heatmap in Figure 3B compared to Figure 3A. Similarly, there were some pairwise estimates of *r*^2^ that were larger under GUS-LD (Figure 4B) compared to the standard likelihood approach (Figure 4A), which is seen by the fact that some of the yellow squares in Figure 4A appear more orange in Figure 4B. The average value of all the pairwise estimates for the two LD measures was larger under GUS-LD than the standard likelihood approach (Table 3). Compared to the simulation results, the difference in the LD estimates between the two approaches is not particularly large. This is due to the fact that there were a number of SNPs with high mean read depths (Figure S5). Nevertheless, this example demonstrates that GUS-LD increases the estimated level of LD compared with the standard estimation approach when there are genotypes with associated low read depths present in the dataset.

**Figure 3.**
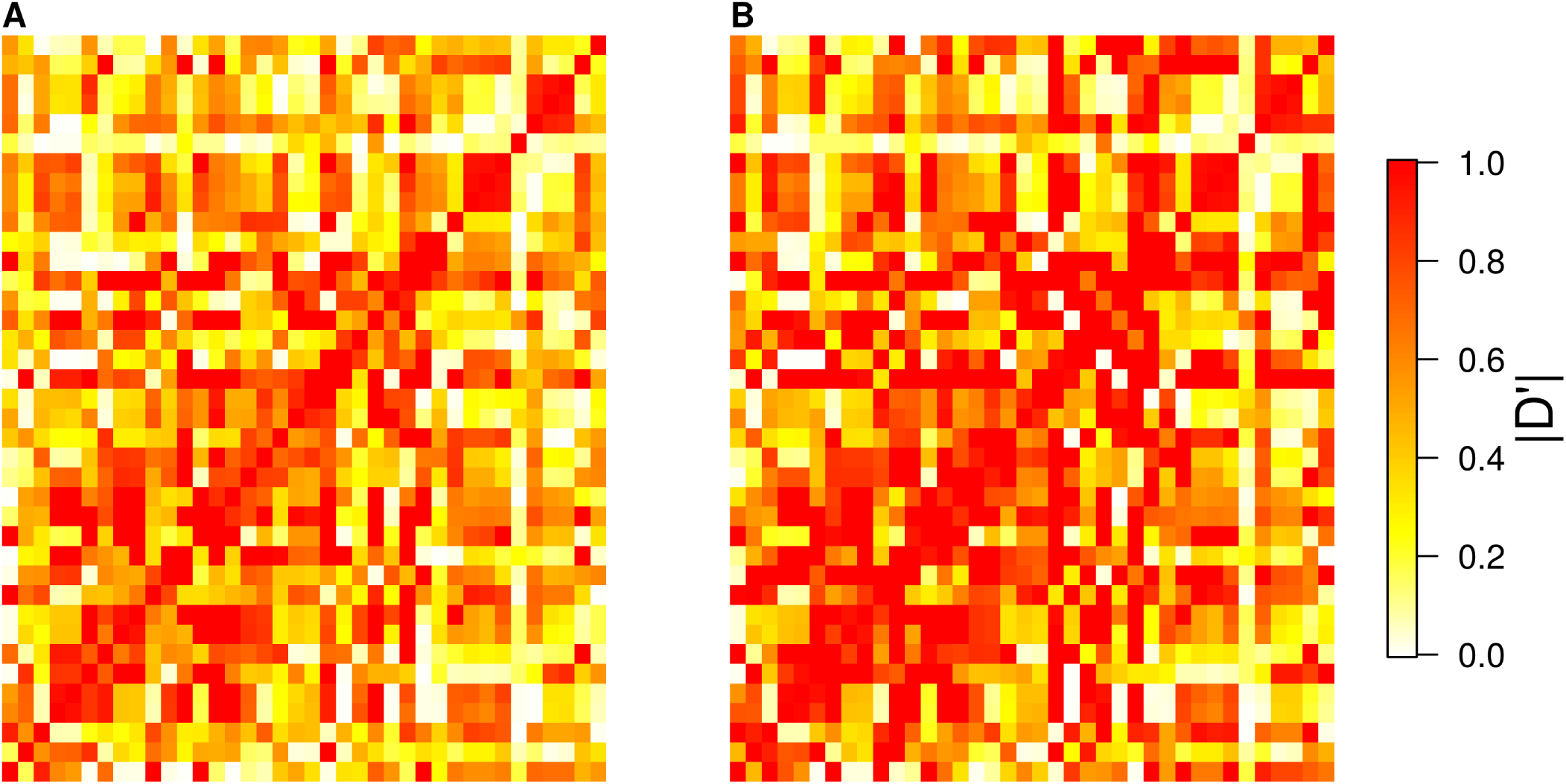
Heatmaps of the absolute value of the pairwise estimates for *D* ^′^ between all 38 SNPs in the deer dataset using (A) the standard likelihood approach which does not account for undercalled heterozygous genotypes and (B) GUS-LD.

**Figure 4.**
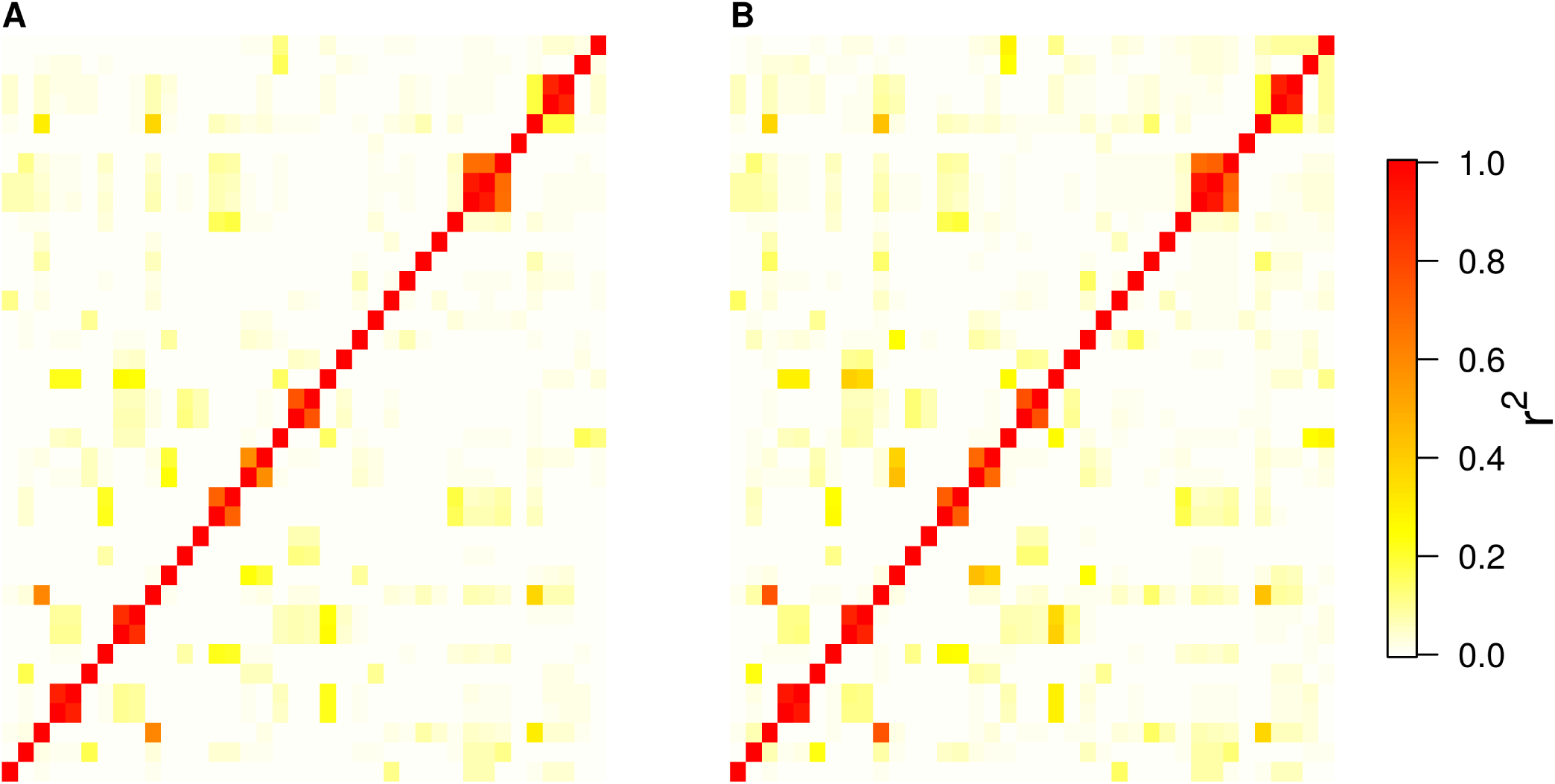
Heatmaps of the pairwise estimates for *r*^2^ between all 38 SNPs in the deer dataset using (A) the standard likelihood approach which does not account for under-called heterozygous genotypes and (B) GUS-LD.

**Table 3.**
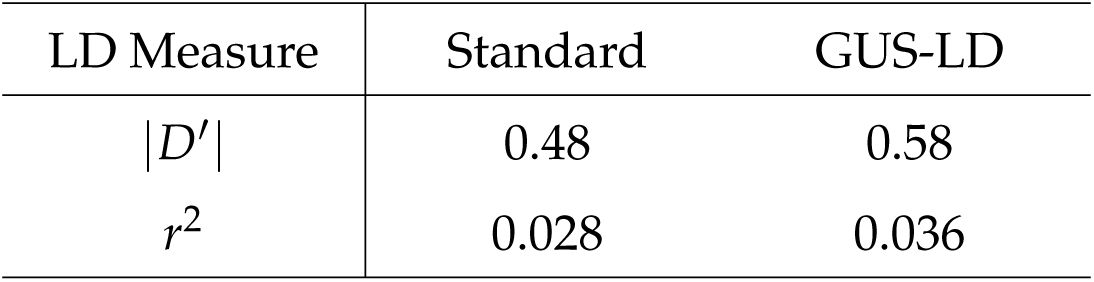
Average LD estimate across all pairs of SNPs for the deer dataset

## Discussion

The introduction of high-throughput sequencing methods that multiplex a large number of individuals is driving forward research into many species, particularly non-model species, and is increasingly being utilized by many researchers. However, analyzing sequencing data using existing analytical tools and methods may, in some cases, be impractical or lead to erroneous results due to the added complexity and nuances of the data compared to other genetic data types. Consequently, the development of new methodological tools for analyzing sequencing data is needed, although the progress of such tools has been slow compared to the sequencing technology (Gardner *et al.* 2014).

Our simulation results have demonstrated that genotyping errors associated with under-called heterozygotes (e.g., allelic dropout) leads to under-estimation of LD, when these errors are not taken into account. This is important as biased estimates of LD can have a profound effect on downstream analyzes. For example, in case-control association studies, it has been shown using simulations that the presence of genotyping errors leads to reduced power in detecting an association between a locus and phenotype (Gordon and Ott 2001; Gordon *et al.* 2002). Russell and Fewster (2009) have also shown via simulations that allelic dropout results in positively biased estimates of effective population size when calculated using LD information. This problem is exacerbated for low coverage data as the rate of genotyping errors is much higher than those used in these simulations studies. We have developed a new method, called GUS-LD, that accounts for under-called heterozygotes in the estimation of LD. Our results show that GUS-LD was able to greatly reduce the bias in the LD estimates at low sequencing depth, although the variability of these estimates were larger compared to the standard approach at low depths, which reflects the additional variation introduced into the data by the uncertainty in the genotype calls at low read depths.

With low coverage sequencing data, there are issues with estimating LD when the true parameter value lies near or on the upper or lower bound of its parameter space (Equations(2)). Specifically, the bias in the LD estimates increases as *D* approaches its upper or lower bound. This is even the case for GUS-LD, which adjusts for genotyping errors associated with low read depths, although the bias is significantly less than the standard likelihood approach. This bias is caused by sampling variation resulting in the maximum of the likelihood (6) lying outside the parameter space of *D*, whereas maximization is performed with respect to the constraint (2). When genotype calls are accurate and without error, this bias, in estimating *D* when its true value is near its upper or lower bound, is absent.

For the methodology developed in this paper, a number of assumptions have been made. Errors resulting from a base being incorrectly called (referred to here as sequencing error) are assumed to be absent. With sequencing errors, a homozygous genotype can be erroneously called as heterozygous or as the opposite homozygote. Neither of these cases can occur as a consequence of low read depth. One approach for estimating sequencing error in sequence data has been proposed by Maruki and Lynch (2014) but requires that addition erroneous alleles not present at that loci are called in the alignment process. In practice, most variant callers by default only allow for two alleles to be called at a SNP, so that estimating sequencing errors using this approach would not be feasible. For GUS-LD, an additional parameter for sequencing error could be included in the model, but the difficulty is in estimating these parameters when the true sequencing error rate is unknown. Secondly, it is assumed that the genotype calls observed in the sequencing data are conditionally independent between loci given the true genotype call. This assumption is reasonable provided that the SNPs are not located on the same sequencing read across the individuals. Estimation of LD is unaffected by the presence of genotyping errors resulting from low read depth when the loci are located on the same read as the true underlying haplotypes in the individuals are preserved. Depending on their settings, many variant callers allow for multiple SNPs to be called on the same sequencing read. However, it is more practical to only retain a single SNP from a given read as the loss of information is minimal and is outweighed by the reduced computational time. Other assumptions include that missing genotypes resulting from read depths of zero occur randomly, and that the alleles of the true genotype are sampled randomly in the sequencing process. If the latter assumption does not hold, one allele will be sampled more frequently than the other (e.g., preferential sampling). In this case, the proportion of heterozygotes seen as homozygotes will be larger than expected under the model, which would result in some bias in the LD estimates at low sequencing depth. If additional information is available, then the probabilities in Equations (5a)-(5e) can be adjusted to reflect alternative sampling models.

The main contributions of this paper are two fold. Firstly, we have demonstrated that there can be significant bias in LD estimates from sequencing data when the read depth is low and the associated errors are not taken into account. This highlights the need for practitioners to either remove these errors by filtering or adjust their methodology to account for these errors. This is particularly important as some LD analyzes give no explicit mention of a minimum cut-off with respect to read depth being used. Secondly, we have proposed GUS-LD as a new method to estimate LD using low coverage sequencing data. GUS-LD will prove valuable to researchers seeking to undertake population studies when cost constraints prohibit the production of high coverage sequencing data or other types of genetic data. In fact, our simulation results suggest that it is more cost-efficient to use low coverage data, as it allows more individuals to be sequenced for the same cost and results in smaller mean square errors for the LD estimates. From our results, the optimal sequencing depth was between 2 and 5, which was similar to the optimal read depth observed by Dodds *et al.* (2015) in the context of relatedness estimation. GUS-LD also allows LD estimation using loci with a mixture of high and low mean read depths, which is particularly useful as the sequencing depth typically varies substantially between SNPs.

## Acknowledgements

This work was funded by FarmIQ (Ministry for Primary Industries’ Primary Growth Partnership fund) – FIQ Systems – Plate to Pasture (PGP06-09020) and the Ministry of Business, Innovation and Employment (New Zealand), Contract C10X1306, “Genomics for Production & Security in a Biological Economy” to AgResearch Ltd. We thank Landcorp Farming Limited for use of their data.

## Footnotes

1 AgResearch, Invermay Agricultural Centre, Private Bag 50034, Mosgiel 9053, New Zealand.

